# Competition constrains parasite adaptation to thermal heterogeneity

**DOI:** 10.1101/2025.09.28.679032

**Authors:** Samuel TE Greenrod, Daniel Cazares, Weronika Slesak, Tobias E Hector, R. Craig MacLean, Kayla C King

## Abstract

Temporal thermal heterogeneity is expected to favour intermediate, generalist phenotypes that can maintain growth across a broad thermal range but have sub-optimal growth at any single temperature. Yet, thermal variation typically occurs in the presence of additional selection pressures which may interact to constrain adaptation to temperature. We propagated competing lytic viral parasites (bacteriophages ϕ14-1 and ϕLUZ19) of *Pseudomonas aeruginosa* under fluctuating temperatures (37-42°C) in monoculture and in co-culture. Without competition, fluctuating temperatures favoured intermediate thermal phenotypes in the phage ϕ14-1 and resulted in more variable evolutionary outcomes compared to static conditions. However, co-selection from fluctuating temperatures and competition led to restricted thermal adaptation, slower evolutionary rates, and fewer putative adaptive mutations in the ϕLUZ19 competitor. Our study highlights the potential for reduced adaptive capacity in interacting communities amidst global climate change.

## Introduction

Thermal heterogeneity plays a key role in shaping species’ evolutionary trajectories. Spanning a broad range of timescales, temperatures fluctuate across multi-year periods (ENSO), between seasons, and even on the order of hours through diurnal (24-hour) cycles. Very slow or rapid thermal fluctuation frequencies, with respect to generation times, typically lead to similar adaptation to static environments through selective sweeps by specialist variants [1,2]. Moderate fluctuation frequencies select for thermal generalists which have intermediate phenotypes across temperatures [1–3]. Generalist phenotypes often arise through the acquisition of multiple specialist mutations [3] or single pleiotropic mutations [4]. Thermal heterogeneity can also promote diversifying selection [5–7] leading to the maintenance of thermal specialist sub-populations [6]. The mechanisms of adaptation to thermal heterogeneity depend on the fluctuation frequency relative to generation time [8]; fluctuations that far exceed generation times in fast-replicating species may favour specialists, but in slow-replicating species may instead select for generalists.

Thermal heterogeneity typically occurs in the context of multiple selective pressures. For example, warming can impose selection on species that are simultaneously adapting to other abiotic stressors or to interactions with predators, competitors, or antagonists [9,10]. The presence of multiple selection pressures can constrain evolution rates through combined negative effects on species fitness which reduce population sizes and mutational supply [11,12]. Co-selection can also restrict adaptation through pleiotropic fitness trade-offs; high fitness under one stressor reduces fitness under another [13,14]. Temporal thermal heterogeneity is expected to promote genetic diversification by increasing niche differences [16] and so may offset the diversity-suppressing impacts of co-selection. However, some studies have indicated that co-selection involving temporal heterogeneity can exacerbate evolutionary constraint [17–19]. The ability of species to adapt to thermal heterogeneity amidst other selection pressures plays an important role in the maintenance of global biodiversity and species extinction risk [11,20–22].

Parasites provide an ideal group of organisms to study adaptation to thermal heterogeneity. Parasites are often exposed to diverse environments and stressors across their multi-stage life cycles. They can have both free-living, vector-based, and host-associated life stages [23]. By moving through numerous external environments during and between replicative cycles, parasites experience high temporal thermal heterogeneity (Greenrod et al., in press; [24]). During the infection stage, parasites can also induce fevers in hosts, driving thermal changes [25]. Finally, parasites are expected to face increasingly frequent thermal extremes as a result of global climate change [26]. While contending with variable thermal environments, parasites must adapt to host immune responses [27] and competition with co-infecting parasites in the same host population or individual [28]. Within and between-host competition are primary determinants of parasite virulence [29] signifying that interactions between competition and environment-based selection can shape parasite evolution [30].

We predicted that thermal heterogeneity would select for generalist parasite populations, which have intermediate phenotypes, and promote genetic diversity [1]. We also predicted that co-selection with other environmental stressors would constrain parasite adaptation [17]. We passaged two lytic viral parasites (thermal generalist ϕLUZ19 and specialist ϕ14-1) under a fluctuating thermal regime (37-42°C) in the absence and presence of a phage competitor. Phages evolved with a static bacterial host, *Pseudomonas aeruginosa*. We compared populations evolved under fluctuating temperatures concurrently with those evolved under a static regime (37°C and 42°C), the latter presented in ref. [31]. We evaluated phage phenotypic adaptation through growth assays at 37°C and 42°C. We also conducted phage population sequencing to identify adaptive mutations and measure evolutionary rates.

## Methods and Materials

### Strains, Storage, and Culture Conditions

This study builds on a previously published experimental framework [31] using the same bacterial host and bacteriophage strains. *Pseudomonas aeruginosa* PAO1 was used as the non-evolving bacterial host throughout. Two lytic phages, ϕLUZ19 and ϕ14-1, were used due to their known thermal response differences: ϕLUZ19 performs well at both 37°C and 42°C, while ϕ14-1 is growth-restricted at 42°C [31,32]. Phage lysates and bacterial stocks were prepared as in refs [31,32].

### Experimental Evolution

The experimental evolution design closely followed that of ref. [31] with additional treatments incorporating fluctuating temperatures. Phages were serially passaged for 15 days under four conditions: monoculture and co-culture, each at either static or fluctuating temperatures (daily shifts between 37°C and 42°C). Each treatment included six independent replicate populations initiated from a single ancestral lysate.

Phages were propagated without shaking with a non-evolving ancestral PAO1 bacterial host. For the initial passage, ancestral phage lysates were diluted to 10^8^ PFU/ml and 300μl were added to 2.7ml 10^8^ CFU/ml bacterial culture in loose-lid 14ml falcon tubes. Phage co-culture populations were prepared by combining 150μl each of ϕLUZ19 and ϕ14-1 10^8^ PFU/ml lysates prior to mixing with bacteria. The initial passage phage densities were ~10^7^ PFU/ml resulting in a phage/bacteria ratio (multiplicity of infection, MOI) = ~0.1. Following addition of bacterial cultures, tubes were incubated statically at 37°C or 42°C in circulating water baths for 8h. Fluctuating passages started and ended at 37°C.

After incubation, phage populations were harvested by centrifugation (3,095×g, 5 min) to pellet bacteria, followed by sterile filtration through 0.2μm filters. Filtrates were stored at 4°C. In subsequent passages, 300μl of lysate was transferred into fresh PAO1 cultures.

### Phage Quantification

Phage titres were determined via the double-layer overlay method [33] following the same protocols as in refs [31,32]. Briefly, bacterial lawns were prepared by mixing 10mL of melted LB-top agar with 300µL of a *P. aeruginosa* PAO1 overnight culture. Phage lysates were serially diluted, and 10µL was spotted onto the bacterial lawns. After incubating plates for 6–8 h at 37°C, spots with the highest number of discernible plaques were counted and reported. ϕLUZ19- or ϕ14-1-resistant PAO1 strains were used for selective plating enabling separate counting of ϕLUZ19 and ϕ14-1 densities in co-cultures. These resistant strains were derived by isolating colonies growing on high titre phage plaques and confirmed via sequencing [31]. All monoculture and co-culture samples were quantified using the appropriate resistant strains to ensure consistency.

### Phage Separation and Concentration

To generate high-titre and pure phage lysates for downstream assays and sequencing, we employed selective double-layer overlays with resistant hosts. Briefly, phages and ϕLUZ19- or ϕ14-1-resistant PAO1 strains were seeded into top agar plates to allow phage propagation. Phages were extracted from plates by scraping top-agar into 15ml falcon tubes containing 5ml of phage buffer (NaCl (100 mM), MgSO_4_ (10 mM), CaCl_2_ (5 mM), Tris-HCl (pH 8) (50 mM), Gelatin (0.01%)). Tubes were mixed overnight after which phages were separated from top agar using sterile-filtration. This process was performed three times to ensure removal of phage competitors from co-culture populations. The purification and extraction protocols were identical to those described in ref. [31].

### Phage Growth Rate Assays

The thermal phenotypes of purified evolved phage populations relative to the ancestor was assessed by measuring phage and bacterial growth across an 8h window under static incubation at 37°C and 42°C. Phage lysates were diluted to 10^5^ PFU/ml and 300uL was mixed with 2.7ml of 10^8^ CFU/ml wild-type PAO1 to a final MOI = ~0.0001. ϕLUZ19 was sampled at 2h, 4h, and 8h; ϕ14-1 at 4h and 8h due to delayed replication. Phage quantification was performed through sterile-filtration through 0.22μm filter plates (Agilent) followed by centrifugation at 2,230xg for 5 mins before spotting onto resistant PAO1 double-layer overlay plates. Each growth rate assay included a single replicate of each evolved phage population and three replicates of the phage ancestor. Growth rate assays were repeated three times across a two-week period to produce three technical replicates.

### Phage Population Genomics

#### DNA Extraction and Sequencing

Phage DNA was extracted from purified lysates as described in [31]. Briefly, ancestral and evolved phage lysates were treated with DNase and RNase to remove bacterial DNA and RNA. Phage particles were lysed using lysis (AL) buffer and proteinase K. Cell debris was precipitated using precipitation (N4) buffer and removed. Finally, DNA was precipitated and washed using isopropanol and ethanol. DNA quality was assessed with NanoDrop 2000c (Thermo Scientific) and quantified with Qubit 4 (Thermofisher). Short-read Illumina sequencing was performed by AZENTA/GENEWIZ using their Microbe-EZ pipeline for evolved and ancestral populations. Bacterial genomes (wild-type and phage-resistant strains) were sequenced by MicrobesNG using long-read approaches.

#### Sequence Analysis

Phage reads were pre-processed with Trim Galore (v.0.5.0) (https://github.com/FelixKrueger/TrimGalore) and downsampled using bbnorm from the bbmap package (v.39.18) (https://sourceforge.net/projects/bbmap/). Reads were then mapped to de novo ancestral assemblies generated with shovill (v1.1.0) (https://github.com/tseemann/shovill) using Bowtie2 (v.2.3.4.2) [34]. Variants were identified using breseq (v.0.36.1) [35]. Ancestral assemblies were annotated with prokka (v.1.14.5) [36], guided by the NCBI GenBank file for each phage (ϕ14-1: NC_011703; ϕLUZ19: NC_010326).

Wild-type and resistant PAO1 genomes were assembled using Autocycler (v. 0.4.0) [37] and polished via Polypolish (v. 0.6.0) [38]. Final assemblies were re-oriented with Dnaapler (v. 1.2.0) [39] and annotated using prokka (v.1.14.5) [36]. The workflow was deployed using a Dockerised Nextflow pipeline (v. 1.0.2) available at https://doi.org/10.5281/zenodo.15706447. Mutations in resistant PAO1 strains were identified by mapping long reads to the wild-type assembly with minimap2 (v.2.24) [40] and variant calling with medaka (v.2.1) (https://github.com/nanoporetech/medaka). All bioinformatic analyses were conducted with default parameters.

### Statistical Analyses and Visualisation

All statistical analyses and data visualisation were conducted using packages in R (v.4.3.2) and RStudio [41,42]. Data wrangling was performed using “Tidyverse” (v.2.0.0) R packages [43]. Phage growth and evolution rates were compared between evolution treatments using linear mixed effect models with the “lme4” (v.1.1-36) R package [44] where the response variable was phage density (pfu/ml) or genetic distance from ancestor, the explanatory variables were an interaction term between evolution treatment and temperature, and batch was a random effect. Within-group variation in genetic distance from ancestor was analysed using Levene’s test. The prevalence of unique compared to shared mutations across evolution treatments was analysed using Fisher’s exact test. Phage genetic distance between groups was also compared by constructing neighbour-joining trees based on Euclidean genetic distance using the “ggtree” (v.3.10.1) R package [45]. Data and code used in analyses can be found at https://github.com/SamuelGreenrod/Evol_fluctuating.

## Results

### Fluctuating temperatures select for generalist phenotypes in monoculture

Fluctuating environments can favour generalists with intermediate phenotypes across conditions [1]. Given ϕ14-1 has previously been shown to grow poorly at 42°C, we hypothesised that ϕ14-1 populations passaged under fluctuating conditions would rapidly adapt to 42°C but have lower fitness at 37°C and 42°C compared to static evolved populations. In monoculture, ϕ14-1 densities increased during 37°C passages but decreased in 42°C passages (Fig. 1A). As phage lysates were diluted 10-fold in between passages, phage density decreases reflect lower than 10-fold ϕ14-1 population growth during 42°C passages. ϕLUZ19 monoculture populations reached and then maintained high densities in all passages. This phage showed low variation in inter-passage densities.

**Figure 1.**
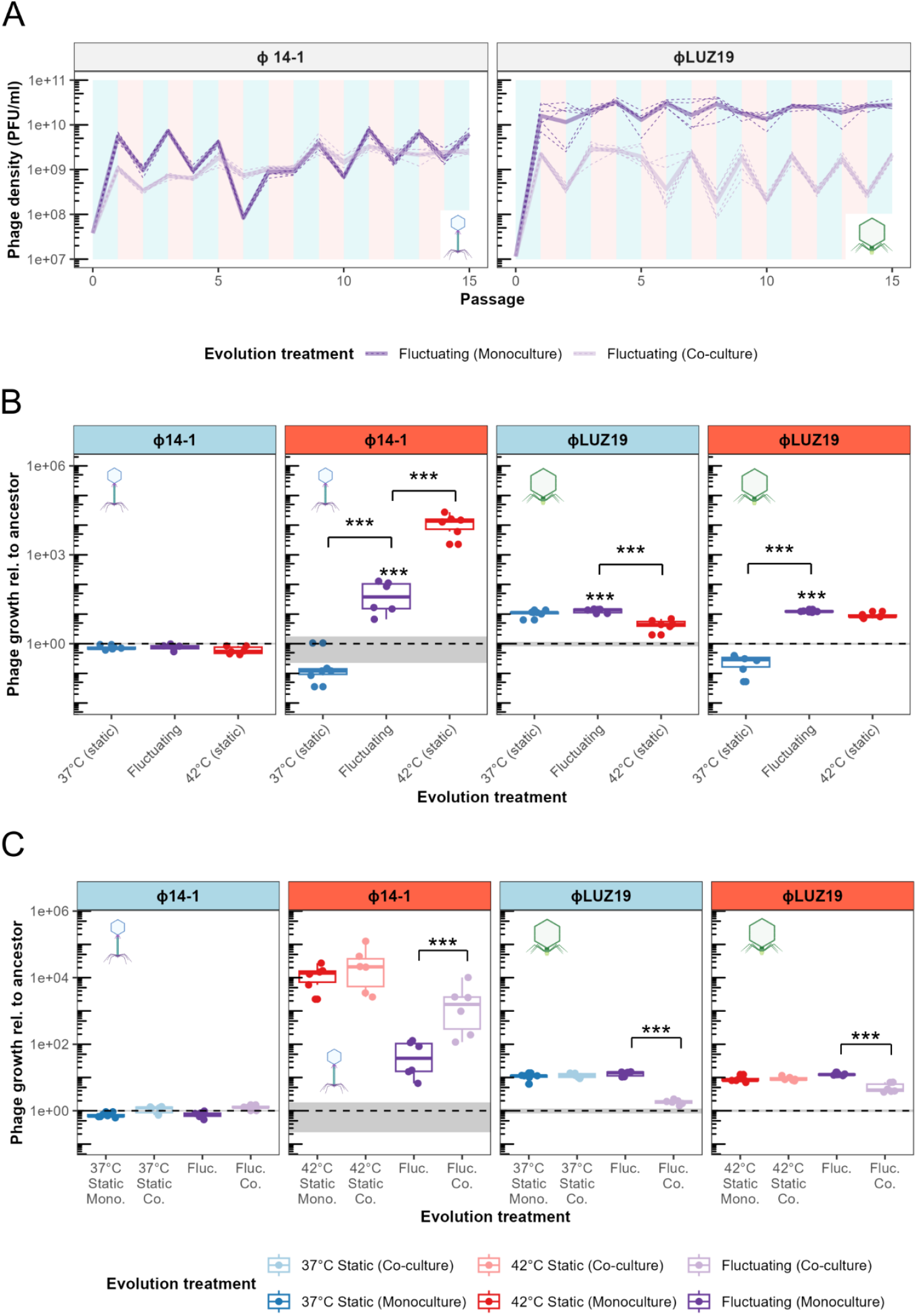
Co-selection in communities constrains adaptation to thermal fluctuations. **A**) Population dynamics of phages passaged in monoculture and co-culture under fluctuating temperatures. Values show densities at the end of each passage prior to dilution. As phage lysates were diluted 10-fold in between passages, density decreases reflect less than 10-fold population growth during passages. Plot background colour reflects the temperature during that passage where light blue is 37°C and light red is 42°C. Phage icons illustrate the two different phages used in the experiments (ϕ14-1, myovirus in blue; ϕLUZ19, autographivirus in green) [46] and are used hereafter to refer to phages in figures. **B**) Fluctuating and static temperature evolved population growth rates relative to the ancestor. Growth rates were measured after 2h for ϕLUZ19 and 4h for ϕ14-1. Six biological replicates were assayed, and data points show the average of three technical replicates. Panel strip colour reflects the temperature that growth was tested at where light blue is 37°C and light red is 42°C. Fluctuating populations are presented in purple with 37°C static populations in blue and 42°C static populations in red. Ancestral growth is shown by dashed grey line with standard errors shown as a grey box (n = 3). *** = p < 0.001. Absence of asterisk reflects non-significance. Static monoculture temperature data was adapted from ref. [31]. **C**) Growth rates of fluctuating and static temperature monoculture evolved populations compared to co-culture evolved populations. Boxes are coloured by evolution treatment with monoculture in dark (37°C static in blue, 42°C static in red, and fluctuating in purple) and co-culture in light (37°C static in light blue, 42°C static in light red, and fluctuating in light purple). Assay temperature and significance values are presented as in Fig. 1B. Six biological replicates were assayed and data points show the average of three technical replicates. Ancestral growth rates and significance signs are presented as in Fig. 1B. Static co-culture data was adapted from ref. [31].

We assessed phage evolution by measuring the growth rates of static and fluctuating evolved populations relative to the ancestral phage through growth assays at 37°C and 42°C (Fig. 1B). We found that growth rates of both phages in monoculture depended on the interaction between evolution treatment and assay temperature (ϕ14-1: F_3,29_ = 125.8, p < 0.001; ϕLUZ19: F_3,29_ = 96.0, p < 0.001). At 42°C, ϕ14-1 fluctuating populations were found to have an intermediate phenotype between those evolved under static conditions. ϕ14-1 fluctuating populations had significantly higher growth rates at 42°C than 37°C static populations (t(29) = −12.4, p < 0.001) but lower growth rates than 42°C static populations (t(29) = 12.7, p < 0.001). At 37°C, ϕ14-1 fluctuating populations had no significant difference to static populations possibly due to phage growth being measured after phages had reached carrying capacity (see ref. [31]). For ϕLUZ19, fluctuating evolved populations had significantly higher growth at 42°C than 37°C static populations (t(29) = −19.3, p < 0.001). However, growth was not significantly different to 42°C evolved populations (t(29) =-1.65, p = 0.37). The opposite findings were observed at 37°C; fluctuating evolved populations had significantly higher growth rates than those evolved at 42°C but similar growth rates to 37°C evolved populations (42°C static: t(29) = −5.2, p < 0.001; 37°C static: t(29) = −0.99, p = 0.75).

### Co-selection from fluctuating temperatures and competition constrains thermal adaptation

The presence of additional selection pressures is expected to constrain adaptation to fluctuating temperatures by reducing mutational supply and compounding fitness trade-offs [11,18,19]. We hypothesised that phages evolved under co-selection from fluctuating temperatures and competition would have lower growth rates at 37°C and 42°C than those evolved under static temperatures or fluctuating monoculture conditions. While ϕ14-1 densities fluctuated between passages in monoculture, co-culture densities rapidly increased and then stabilised between passages (Fig. 1A). In contrast, ϕLUZ19 populations were stable in monoculture, but during fluctuations in co-culture, experienced high growth at 37°C and low growth at 42°C.

We then assessed evolved phage growth rates at 37°C and 42°C (Fig. 1C). We found a significant interaction between evolution treatment (monoculture and co-culture) and temperature regarding growth rates for both phages (ϕ14-1: F_6,59_ = 75.0, p < 0.0001; ϕLUZ19: F_6,59_ = 103.5, p < 0.0001). While competition had no impact on the growth rates of static ϕ14-1 populations (37°C: t(59) = −0.78, p = 0.99; 42°C: t(59) = −0.98, p = 0.99), fluctuating co-culture evolved populations had significantly higher growth rates at 42°C compared to populations evolved in monoculture (t(59) = −6.7, p < 0.0001). No significant difference was observed at 37°C (t(59) = −1.0, p = 0.99). Similar to ϕ14-1, there was no impact of competition on static evolved ϕLUZ19 population growth rates (37°C: t(59) = −0.42, p = 1.0; 42°C: t(59) = −0.21, p = 1.0). However, ϕLUZ19 populations evolved with fluctuations and competition had significantly lower growth rates at both 37°C and 42°C compared to monoculture (37°C: t(59) = 10.1, p < 0. 0001; 42°C: t(59) = 4.9, p < 0.001).

### Fluctuating environments favour specialist mutations

Fluctuating temperatures generally select for multiple specialist mutations [3]. We hypothesised that fluctuating evolved populations would show genetic similarities to both high and low temperature static populations. Phage genomic evolution was assessed by constructing neighbour-joining trees of end-point populations based on Euclidean genetic distances (Fig. 2A). Genetic distances were calculated based on the presence and frequency of genetic variants (SNPs, indels) that had > 10% frequency. Fluctuating evolved populations did not form a unique clade but instead were found to co-locate with either high or low temperature static populations. ϕ14-1 fluctuating populations were distributed across the tree and generally did not cluster with static populations. Conversely, ϕLUZ19 fluctuating populations were primarily found within the 42°C static clade.

**Figure 2.**
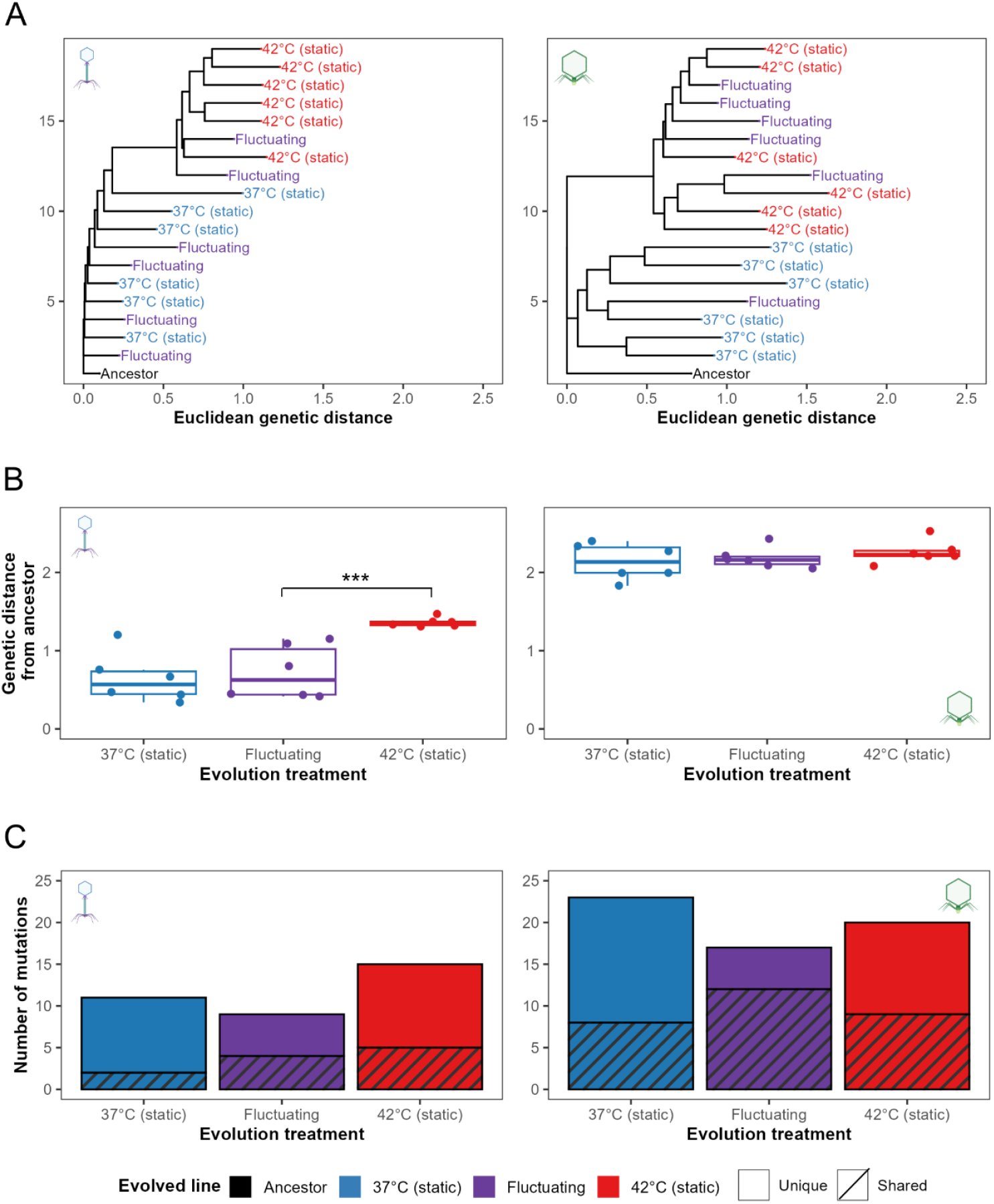
Fluctuating temperatures select for specialist mutations. **A**) Neighbour-joining trees of evolved and ancestral phage populations constructed using Euclidean genetic distances. Genetic distances were calculated based on the presence and frequency of mutations present at > 10% frequency. Tree is rooted at the ancestor and populations are coloured by evolved treatment. **B**) Phage evolution rates, measured based on Euclidean genetic distance from the ancestor, for evolved monoculture phage populations. *** = p < 0.001. **C**) Stacked bar charts show the number of high frequency (>20% frequency), putative adaptive variants that are unique to or shared between evolution treatments. Bars are coloured by evolution treatment. Unique genes are shown as clear bars and shared genes are shown as striped bars. Static temperature data was adapted from ref. [31].

We further analysed static and fluctuating population genetic similarities by measuring evolution rates based on Euclidean genetic distance from ancestor (Fig. 2B). For ϕ14-1, fluctuating evolved populations had significantly lower evolution rates than 42°C static populations (t(15) = −4.1, p < 0.01). However, evolution rates were equal between fluctuating and 37°C static populations (t(15) = −0.51, p = 0.87). There was no significant difference in evolution rates between ϕLUZ19 fluctuating populations and either 37°C or 42°C static populations (37°C: t(15) = −0.46, p = 0.89; 42°C: t(15) = −0.75, p = 0.74). Notably, ϕ14-1 fluctuating populations had significantly greater within-group variation in evolution rates compared to 42°C static populations (t(15) = 2.9, p < 0.05) but not 37°C static populations (t(15) = −0.70, p = 0.77). ϕLUZ19 fluctuating populations had no significant difference in within-group variation compared to static populations (37°C: t(15) = 2.0, p = 0.15; 42°C: t(15) = −0.10, p = 0.99).

We then determined the prevalence of individual genetic variants (SNPs, indels) that were unique to or shared between evolution treatments (Fig. 2C). Only putative adaptive variants with >20% frequency were included. For ϕ14-1, 2/11 (18%) of 37°C static variants and 5/15 (33%) of 42°C static variants were shared with other evolution treatments compared to 4/9 (44%) variants in fluctuating evolved populations. For ϕLUZ19, shared variants constituted 8/32 (35%) of 37°C static and 9/20 (45%) of 42°C static variants compared to 12/17 (71%) in fluctuating evolved populations. To assess the overall impact of evolution treatment on the ratio of unique and shared mutations, we pooled mutations from ϕ14-1 and ϕLUZ19 observing a significant difference in the prevalence of shared mutations relative to unique mutations between evolution treatments (Fisher’s exact test: p < 0.05). Significant differences were not observed when analysing phages independently (ϕ14-1, p = 0.47; ϕLUZ19, p = 0.08). ϕ14-1 fluctuating mutations were primarily shared with 42°C static populations (Fig. S1). In contrast, ϕLUZ19 fluctuating mutations were shared equally with 37°C and 42°C static populations.

Finally, we investigated which mutations drove clustering between fluctuating and static populations (Fig. S2; Table S1). While ϕ14-1 fluctuating and 37°C static populations showed little clustering, two fluctuating populations clustered with the 42°C static clade. These two replicate populations contained parallel deletions in a hypothetical protein with high similarity to a DNA ligase (BlastP: 97.47% identity, 95% sequence overlap with Pseudomonas phage PhL_UNISO_PA-DSM_ph0031 DNA ligase protein), previously identified in all ϕ14-1 42°C static populations [31]. The clustering of 5/6 ϕLUZ19 fluctuating populations with the 42°C static populations were attributed to parallel insertions in an intergenic region between two hypothetical proteins. This intergenic insertion was also previously identified in all ϕLUZ19 42°C static populations [31].

### Co-selection constrains molecular evolution

By reducing growth rates, co-selection from fluctuating temperatures and competition are expected to reduce evolution rates and restrict the acquisition of adaptive mutations [11]. Due to their elevated growth rates, we hypothesised that ϕ14-1 fluctuating co-culture populations would have higher evolution rates than monoculture populations. No significant difference in evolution rate was observed for ϕ14-1 co-culture populations compared to monoculture populations (F_1,10_ = 4.1, p = 0.07) (Fig. 3A). However, non-significance was driven by a single low evolution rate replicate in the co-culture treatment; when the replicate was removed, co-culture populations had significantly greater evolution rates than monoculture (F_1,9_ = 8.6, p < 0.05). We also found no significant difference in within-group variation in evolution rates between monoculture and co-culture populations (t(10) = 1.50, p = 0.17), although the difference was also significant once the low evolution rate co-culture replicate was removed (t(10) = 3.6, p < 0.01).

**Figure 3.**
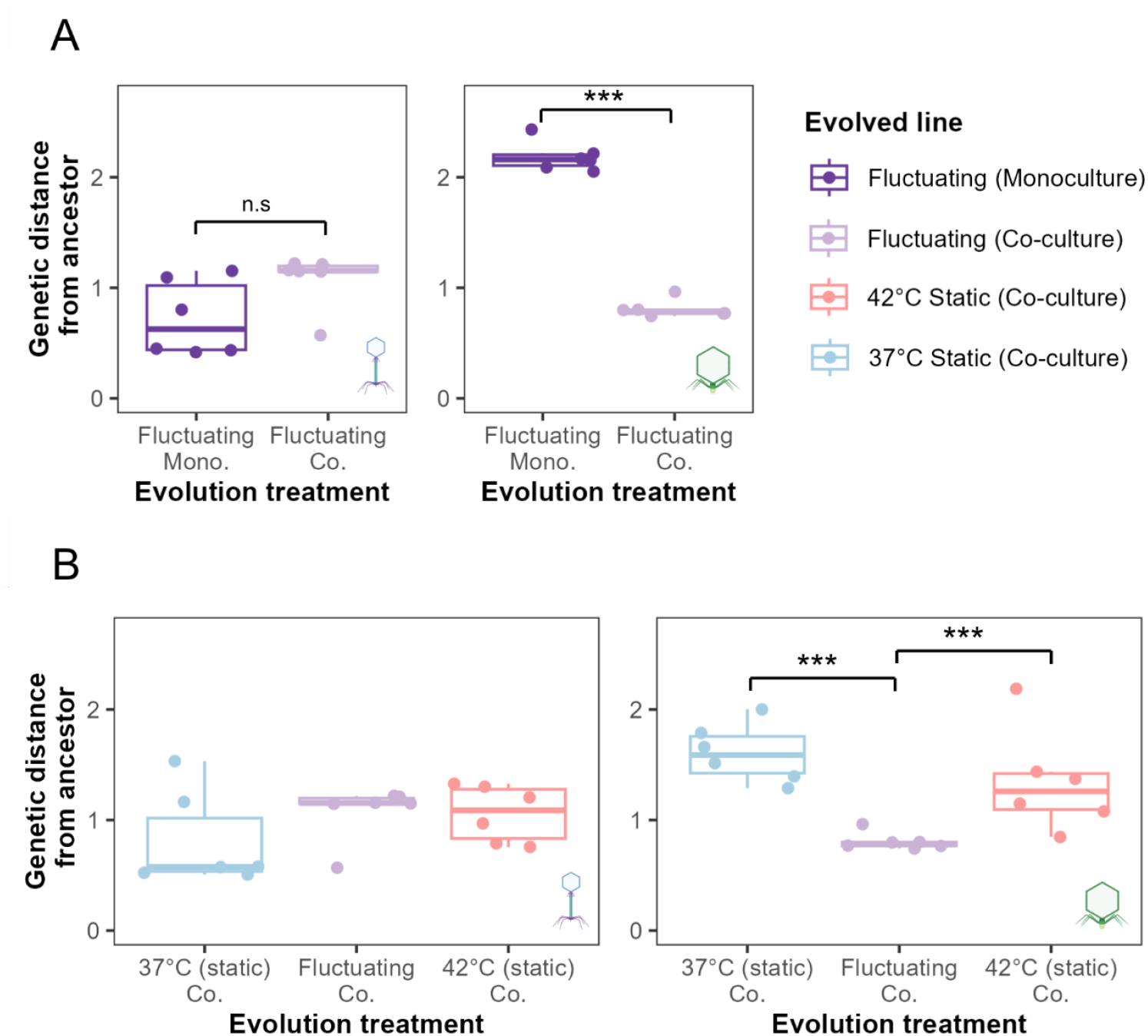
High environmental complexity constrains evolution rates. **A**) Evolution rates, measured based on Euclidean genetic distance from the ancestor, of phage populations evolved under fluctuating temperatures in monoculture (deep purple) and co-culture (light purple). **B**) Evolution rates of 37°C static, fluctuating, and 42°C static co-culture populations. Plot layout is the same as panel A. *** = p < 0.001. N.S is used to denote lack of statistical significance. Static temperature data was adapted from ref. [31].

For ϕLUZ19, we hypothesised that fluctuating co-culture populations would have lower evolution rates than monoculture populations due to the suppression of ϕLUZ19 by ϕ14-1. ϕLUZ19 fluctuating co-culture populations had significantly lower evolution rates than monoculture populations (F_1,10_ = 470, p < 0.001). We further hypothesised that ϕLUZ19 fluctuating co-culture populations, but not ϕ14-1 populations, would have lower evolution rates than 37°C or 42°C static co-culture populations. Evolution rates were equal between co-culture populations for ϕ14-1 (F_2,15_ = 1.2, p = 0.3) (Fig. 3B). However, ϕLUZ19 fluctuating co-culture populations were found to have significantly lower evolution rates than both 37°C static and 42°C static monoculture populations (37°C: t(15) = 4.5, p < 0.01; 42°C: t(30) = 3.0, p < 0.05).

Finally, we assessed the impact of competition on the acquisition of high frequency mutations (> 20% frequency) in fluctuating populations (Fig. S3). For ϕ14-1, while populations no longer acquired singleton mutations, all populations acquired a deletion or SNP in a putative DNA ligase gene. Mutations in this gene are thought to contribute to high temperature adaptation [31] and were also found in the two monoculture populations which clustered with 42°C static populations in Fig. 2A. For ϕLUZ19, while co-culture populations maintained mutations in tail fiber genes, the populations no longer acquired singleton mutations or the intergenic insertion associated with 42°C static populations.

## Discussion

Under fluctuating selection, both phages were found to rapidly evolve increased growth rates at high temperatures. For phage ϕ14-1, fluctuating temperatures favoured intermediate thermal phenotypes with lower growth rates at high temperatures compared to phages evolved under static temperatures. The evolution of intermediate growth rates is likely due to weaker selection for adaptation to high temperatures in the fluctuating treatment.

Alternatively, adaptation to high temperatures may be constrained during fluctuations due to fitness trade-offs at lower temperatures [47]. For phage ϕLUZ19, fluctuating temperature populations had the same growth rates as static-evolved populations when measured at their evolved temperature. Given ϕLUZ19 was shown to exhibit growth rate trade-offs under static selection [31], this finding is indicative of a no-cost generalist strategy [48]. The lack of growth rate costs may reflect an epistatic pleiotropy, whereby the costs of adaptive mutations depend on the genetic background [48]. Costs may also occur in unmeasured traits such as virulence [49] or tolerance to other stresses [14]. These findings highlight that, while fluctuating temperatures select for greater high temperature growth, the exact phenotypic outcomes of fluctuating selection vary between phage taxa.

We found that fluctuating temperatures resulted in more variable evolutionary trajectories; while static evolved populations generally formed clusters, fluctuating populations were genetically similar to both 37°C and 42°C static evolved populations. Further, we found that parallel mutations acquired under fluctuating selection were the same as those previously identified in static evolved populations [31]. These findings could be explained by historical contingency [50] whereby random mutations that are adaptive at low or high temperatures become fixed in a subset of populations [6]. Depending on the timing of mutation appearance, fluctuating populations may then resemble individual static environments. Fluctuating environments have also been shown to select for mutations conferring fitness in the most extreme environment [51]. The clustering of ϕLUZ19 fluctuating populations with 42°C static populations likely reflects selective sweeps combined with asymmetrical selection where 42°C adaptive variants are fixed more rapidly.

While fluctuating environments can promote genetic diversification, co-selection with other environmental stressors can constrain adaptation [11]. Uiterwaal et al [18] showed that combined fluctuating temperatures and predation restricted adaptation to both selection pressures in *Paramecium caudatum* populations. Similar findings have also been observed in *Daphnia magna* with co-selection from thermal fluctuations and predation/pollutants [19,52]. We found that combined fluctuating temperature- and competition-based selection constrained both thermal adaptation and slowed evolutionary rates in phage ϕLUZ19. Further, co-selection resulted in greater ϕLUZ19 evolutionary constraint with fluctuating temperatures than static temperatures. The negative effects of combined environmental stressors are often non-additive and instead exhibit synergies [11]. We have previously shown that selection from high temperatures synergises with competition to constrain ϕLUZ19 evolution rates [31]. The present study extends these findings by showing that the selective synergy between temperature and competition is greater under fluctuating temperatures than static temperatures. One potential explanation for these findings is that fluctuating environments reduce the strength of selection for adaptive mutations [53]. Weaker directional selection combined with suppression by competitors may constrain both the supply and fixation of beneficial mutations leading to particularly low evolution rates.

Anthropogenic activities, including global climate change, mean that species are facing increasingly variable and complex environments [54]. With ongoing global biodiversity loss, species must adapt to tolerate environmental stressors to avoid extinctions. Our findings highlight that while species can rapidly adapt in response to thermal variation, co-selection with other stressors, such as competition, may restrict species adaptive capacity. These results have particular relevance for parasites which must simultaneously adapt to both thermal heterogeneity, community competition in coinfections, and host immune responses. With global parasite biodiversity at risk due to climate change [26], evolutionary constraint caused by co-selection may prevent parasite adaptation to thermal stress and further increase the probability of parasite extinctions. Future studies should consider the evolution-constraining effects of co-selection when assessing species extinction risk in thermally variable environments.

## Acknowledgments

We thank R. Salguero-Gomez, T. Richards, T. Barraclough, and K. Foster for feedback on the experimental design and results. This work was supported by the Biotechnology and Biosciences Research Council (BB/T008784/1) to S.T.E.G. as well as the Natural Environment Research Council (NE/X000540/1) and NSERC Canada Excellence Research Chair to K.C.K. The funders had no role in study design, data collection and interpretation, or the decision to submit the work for publication. Phage sequence reads are accessible on NCBI (https://www.ncbi.nlm.nih.gov/) under BioProject ID: PRJNA1334331.

## Extended data

**Figure S1.**
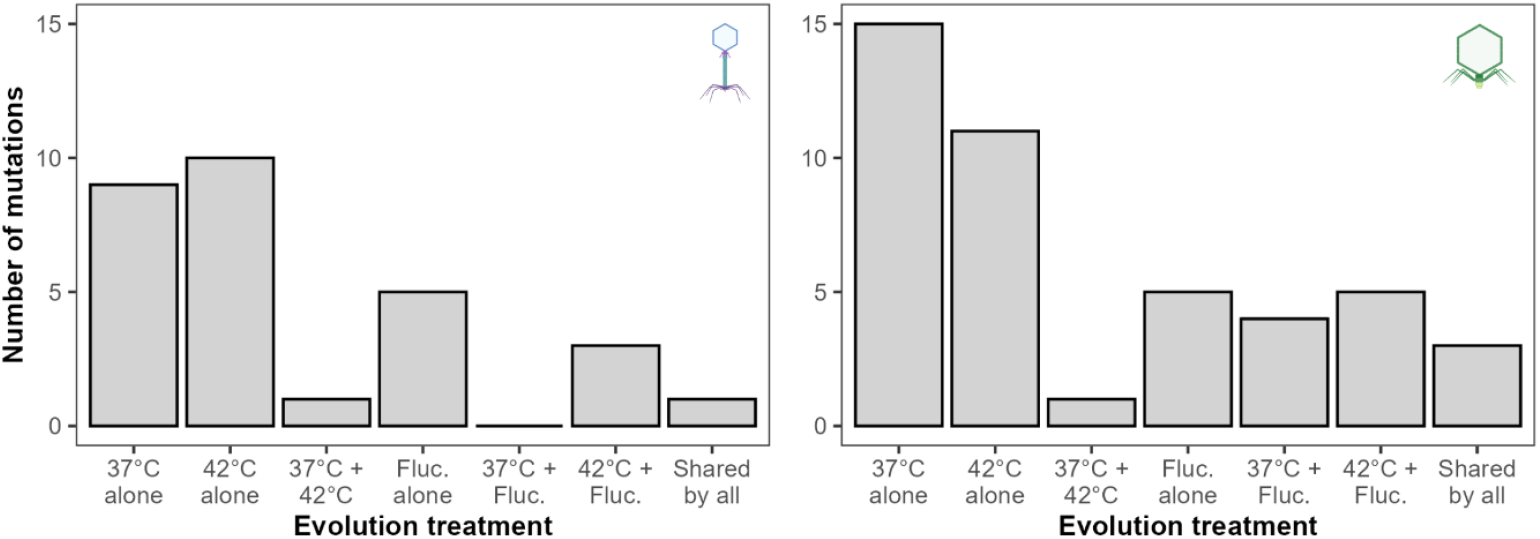
Bar charts show the number of high frequency variants (> 20% frequency) that are unique to or shared between evolution treatments. Variants are shown as either unique to individual evolution treatments, shared between two treatments, or shared by all three. Static temperature data was adapted from ref. [31].

**Figure S2.**
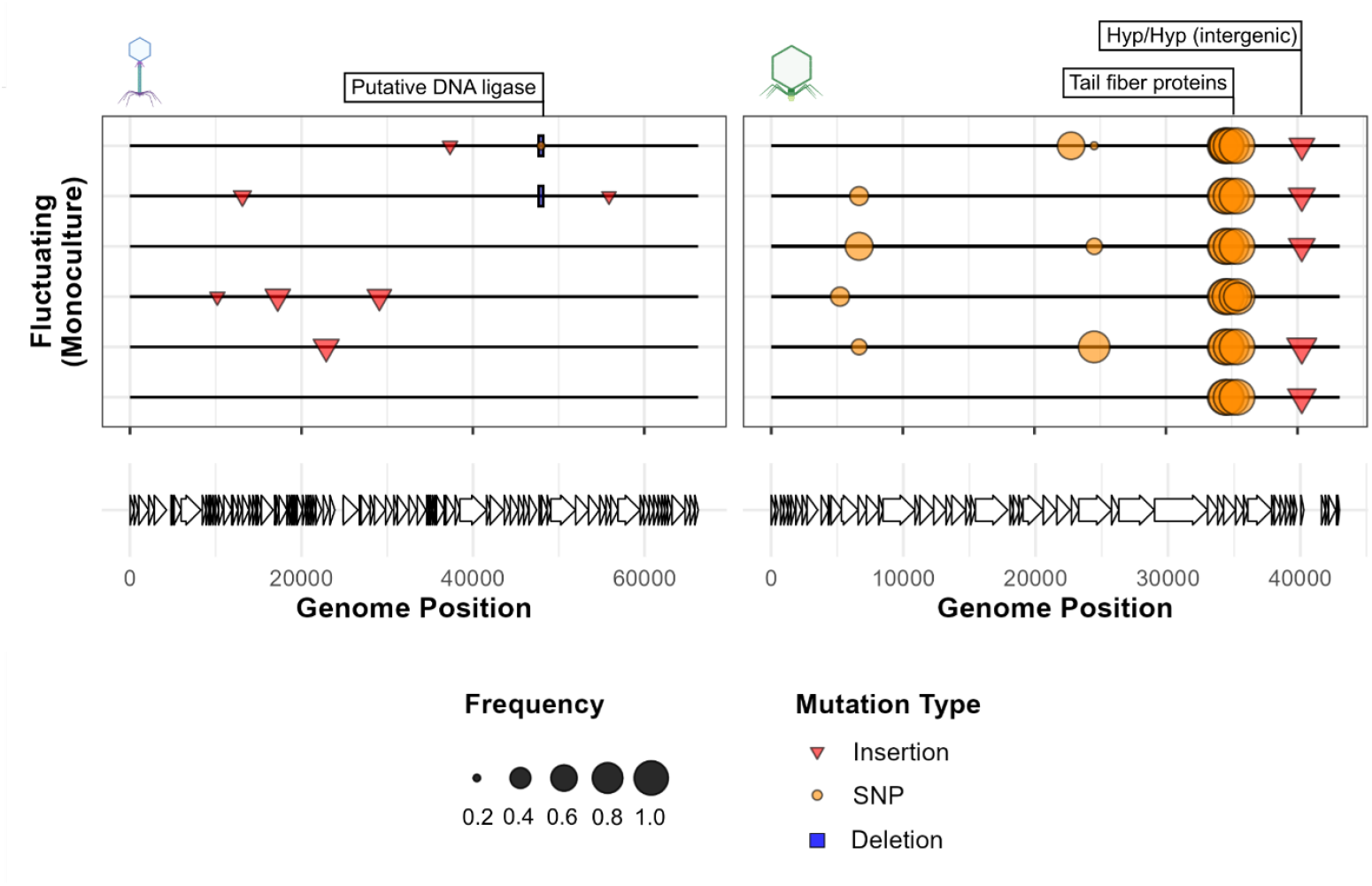
Fluctuating temperatures select for specialist mutations. Mutation plots show genetic variants associated with fluctuating temperatures in monoculture in phage populations. Lines represent individual biological replicates. Symbols within plots show variants across the phage genome at > 20% prevalence and which were not observed in the ancestral population. Length of deletion bars represent the size of deletion except for the ϕ14-1 deletion at ~48kb which is a 1bp deletion but given a fixed size for visibility. Labels show gene annotations for mutations found in 37°C and 42°C evolved populations [31]. Putative DNA ligase in ϕ14-1 was originally annotated a hypothetical protein but has high homology to Pseudomonas phage PhL_UNISO_PA-DSM_ph0031 DNA ligase protein.

**Figure S3.**
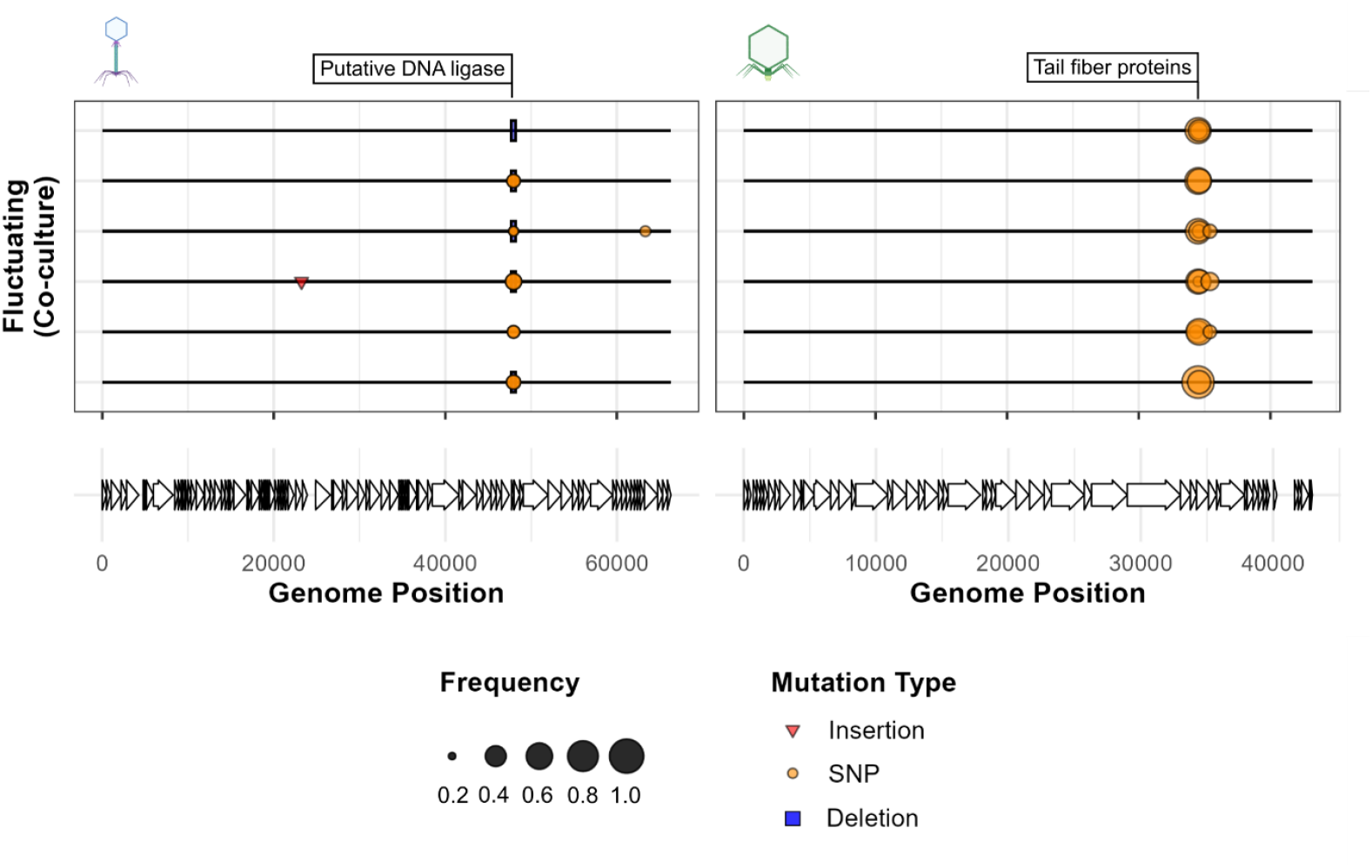
Competition restricts the acquisition of high frequency mutations. Mutation plots show genetic variants associated with fluctuating temperatures in co-culture in phage populations. Lines represent individual biological replicates. Symbols within plots show variants across the phage genome at > 20% prevalence and which were not observed in the ancestral population. Length of deletion bars represent the size of deletion except for the ϕ14-1 deletion at ~48kb which is a 1bp deletion but given a fixed size for visibility. Labels show gene annotations for mutations found in 37°C and 42°C evolved populations [31]. Putative DNA ligase in ϕ14-1 was originally annotated a hypothetical protein but has high homology to Pseudomonas phage PhL_UNISO_PA-DSM_ph0031 DNA ligase protein.

